# CRISPR editing of *sftb-1*/SF3B1 in *C. elegans* allows the identification of synthetic interactions with cancer-related mutations and the chemical inhibition of splicing

**DOI:** 10.1101/634634

**Authors:** Xènia Serrat, Dmytro Kukhtar, Eric Cornes, Anna Esteve-Codina, Helena Benlloch, Germano Cecere, Julián Cerón

**Affiliations:** Modeling human disease in C. elegans Group. Genes, Disease and Therapy Program, Institut d’Investigació Biomèdica de Bellvitge–IDIBELL. 08908, Barcelona, Spain.; Mechanisms of Epigenetic Inheritance Group. Institut Pasteur. 75724, Paris, France.; CNAG-CRG, Centre for Genomic Regulation (CRG), The Barcelona Institute of Science and Technology, Baldiri Reixac 4, Barcelona 08028, Spain; Universitat Pompeu Fabra (UPF), Barcelona, Spain

## Abstract

*SF3B1* is the most frequently mutated splicing factor in cancer. Mutations in *SF3B1* confer growth advantages to cancer cells but they may also confer vulnerabilities that can be therapeutically targeted. In contrast to other animal models, SF3B1 cancer mutations can be maintained in homozygosis in *C. elegans*, allowing synthetic lethal screens with a homogeneous population of animals. These mutations cause alternative splicing (AS) defects in *C. elegans*, as it occurs in *SF3B1*-mutated human cells. In a screen, we identified RNAi of U2 snRNP components that cause synthetic lethality with *sftb-1/SF3B1* mutations. We also detected synthetic interactions between *sftb-1* mutants and cancer-related mutations in *uaf-2/U2AF1* or *rsp-4/SRSF2*, demonstrating that this model can identify interactions between mutations that are mutually exclusive in human tumors. Finally, we have edited an SFTB-1 domain to sensitize *C. elegans* to the splicing inhibitor pladienolide B. Thus, we have established a multicellular model for SF3B1 mutations amenable for high-throughput genetic and chemical screens.

## INTRODUCTION

Deregulated RNA splicing is emerging as a new hallmark of cancer following the discovery of several splicing factor genes harboring somatic mutations in different tumor types (Urbanski et al., 2018). *SF3B1*, a core component of the U2 snRNP, is the most frequently mutated splicing factor in human cancers. Somatic heterozygous mutations in *SF3B1* are particularly prevalent in myelodysplastic syndromes (MDS)—up to 80% in refractory anemia with ring sideroblasts (Malcovati et al., 2015; Papaemmanuil et al., 2011; Yoshida et al., 2011); and in 15% of chronic lymphocytic leukemia (CLL) (Quesada et al., 2011; te Raa et al., 2015; Wang et al., 2011). *SF3B1* mutations have also been reported in solid tumors, including 20% of uveal melanomas (UM) (Furney et al., 2013; Harbour et al., 2013; Martin et al., 2013), 3% of pancreatic ductal adenocarcinomas (Biankin et al., 2012), and 1.8% of breast cancers (Maguire et al., 2015). RNA-sequencing (RNA-seq) analyses in different tumor types and cell lines with *SF3B1* missense mutations, including the most prevalent substitution K700E, identified distinct alternative splicing (AS) defects (Darman et al., 2015; DeBoever et al., 2015; Liberante et al., 2019).

Recently, CRISPR/Cas9 genome editing has allowed researchers to faithfully reproduce human pathological mutations in animal models. Human *SF3B1* mutations have been introduced in cell lines (Alsafadi et al., 2016; Gupta et al., 2019; Liberante et al., 2019; Shiozawa et al., 2018), but mimicking these mutations in multicellular organisms has only been achieved by conditional alleles in murine models (Xu et al., 2019). Taking advantage of the extraordinary conservation of splicing factors across evolution and the ease of genetic manipulation of *C. elegans*, we established a multicellular model to study *SF3B1* cancer-related mutations. Spliceosome components, and particularly *SF3B1*, are being intensively studied as a target of antitumor drugs (Bonnal et al., 2012; DeNicola and Tang, 2019; Effenberger et al., 2017). Some natural compounds inhibit splicing by targeting SF3B1. Among them, pladienolide B (PB) is particularly relevant because H3B-8800, a PB derivative, has antitumoral effects and is in clinical trials (Seiler et al., 2018a). The inhibitory activity of PB is highly dependent on SF3B1 structure, and single amino acid substitutions cause resistance to PB (Teng et al., 2017; Yokoi et al., 2011). Hence, particular residues in the drug binding site that are not conserved in *sftb-1* may confer resistance to PB in worms, as it has been recently observed in yeast (Hansen et al., 2019).We edited four amino acids to humanize the HEAT repeat 15 in SFTB-1, making *C. elegans* sensitive to PB. Thus, we have created the first *C. elegans* strain sensitive to chemical inhibition of splicing.

Altogether, the multicellular model of *SF3B1* mutations described in this study provides a new pre-clinical platform for identifying new targets and small molecules for use in cancer therapies.

## RESULTS

### *sftb-1*, the *C. elegans* ortholog of *SF3B1*, is an ubiquitously expressed gene essential for development

The *C. elegans* protein SFTB-1 is 66% identical to human SF3B1 in terms of amino acid composition. Homology is particularly high at the HEAT domain, reaching 89% identity, and the most frequently mutated amino acids in cancer are conserved. The SFTB-1 sequence also conserves some of the U2AF ligand motifs (ULMs) that bind U2AF homology motifs (UHMs) present in other splicing factors (Figure 1A **and S1**) (Loerch et al., 2018).

**Figure 1.**
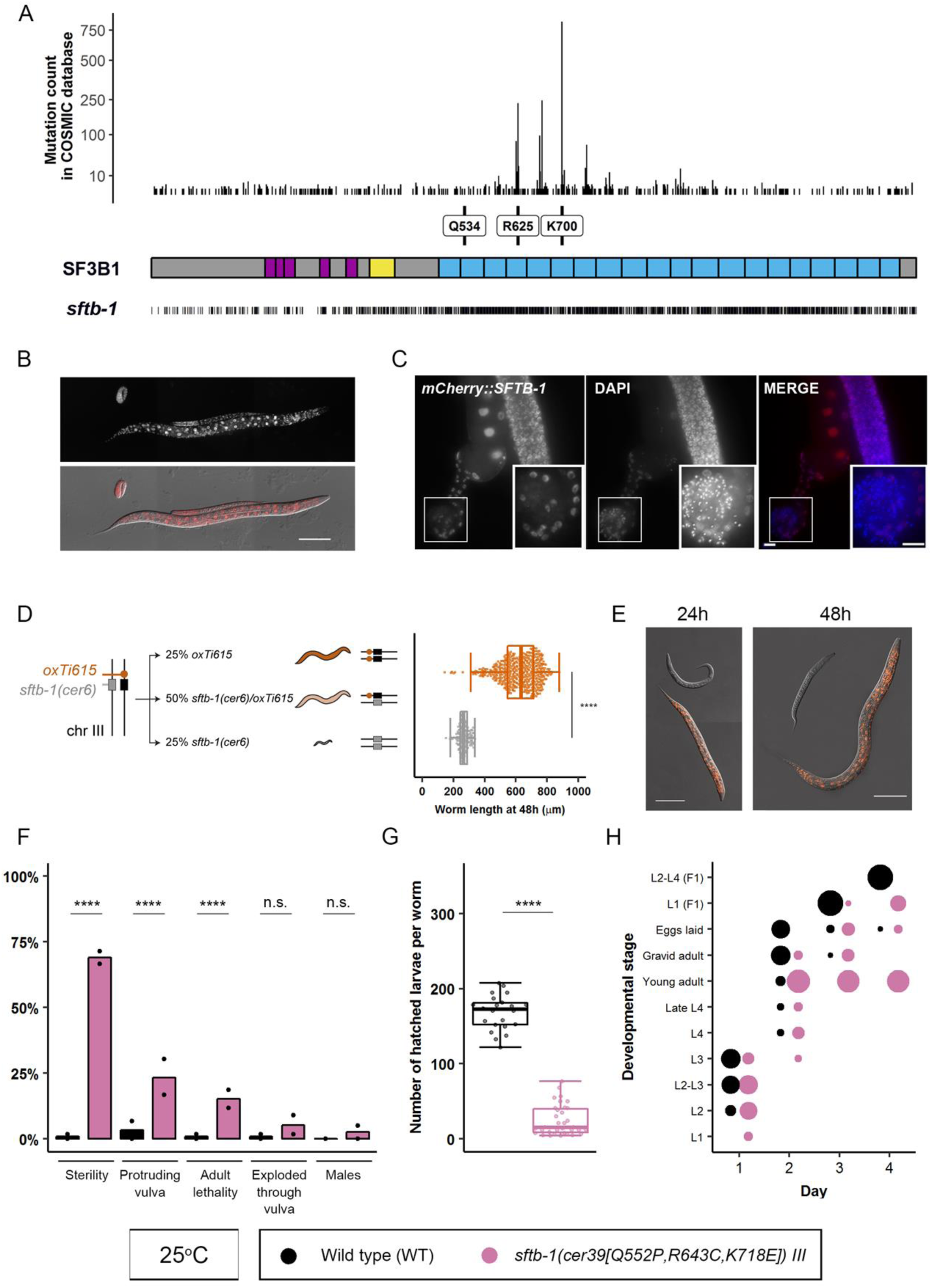
Characterization of *sftb-1*, the *C. elegans* ortholog of *SF3B1*, and different *sftb-1* mutations. (**A**) Top: Number of mutations annotated in COSMIC database (Forbes et al., 2017) at each amino acid position. Middle: schematic structure of the SF3B1 protein. Purple boxes—U2AF ligand motifs (ULMs); yellow box—p14-interacting region; blue boxes—20 HEAT repeats. Labels indicate the three residues that are relevant to this study. Bottom: bars indicate residues that are conserved in the *C. elegans* ortholog *sftb-1*. (**B**) Representative image of an embryo, L2, and L4 larvae showing *mCherry::SFTB-1* endogenous expression. The fluorescence image (top panel) is merged with the corresponding differential interference contrast (DIC) image (bottom panel). (**C**) Fluorescence images of dissected gonads from adult worms expressing *mCherry::SFTB-1*. Gonads were stained with DAPI to visualize nuclei. Insets show differentiated sperm nuclei, where *mCherry::SFTB*-1 is not expressed. (**D**) Left: diagram depicting how the *cer6* deletion allele is maintained in heterozygosis. The WT *sftb-1* copy is genetically linked to a red fluorescent marker (*oxTi615*). Right: worm body length quantification of WT and heterozygotes (red), and *cer6* homozygotes (gray) grown for 48h at 20°C (n=475, 134; N=3). (**E**) Representative images of synchronized red and non-red siblings after growing for 24h and 48h at 20°C. (**F**) Incidence of different *sftb-1[Q552P, R643C, K718E]* phenotypes at 25°C. Dots represent percentages observed in each replicate (n=119, 116; N=2). (**G**) Total number of progeny laid by WT or *sftb-1[Q552P, R643C, K718E]* fertile worms at 25°C (n=22, 35; N=2). (H) *sftb-1[Q552P, R643C, K718E]* mutant displays developmental delay at 25°C. The proportion of the population at each stage is represented by the size of the dots (n=119, 116; N=2). In (D) and (G), dots represent measures in individual worms, and overlaid Tukey-style boxplots represent the median and interquartile range (IQR). Whiskers extend to the minimum and maximum values or at a maximum distance of 1.5xIQR from the box limits. Statistics: (D) and (G), Mann-Whitney’s test; (F), Fisher’s exact test. n.s., no significant difference, **** p< 0.0001. Scale bars: 100 µm (B and E), 10 µm (C). See also **Figures S1, S2 and S3**.

As expected for a core splicing factor, the CRISPR-engineered endogenous fluorescent reporter *mCherry::SFTB-1* showed ubiquitous expression in somatic and germ cells, being absent in mature sperm only (Figure 1b **and** 1c). We also used CRISPR to generate the *sftb-1(cer6)* mutation, a deletion allele that produces a premature stop codon (**Figure S2a**) and causes an arrest at early larval stages, confirming that *sftb-1* is essential for development (Figures 1d **and** 1e). *sftb-1(cer6)* homozygous animals completed embryonic development but arrested as larvae that could undergo few germ cell divisions (**Figure S2b**), suggesting that maternal wild-type (WT) product can still be present in the early larva.

### Different *sftb-1* missense mutations display additive effects when combined

We edited the *C. elegans* genome to mimic the K700E mutation, the most frequent *SF3B1* substitution in tumors. *sftb-1*[*K718E*] homozygous animals did not display any obvious phenotypes. Thus, we decided to reproduce two other *SF3B1* cancer-related mutations. R625C is a particularly prevalent mutation in uveal melanoma, and Q534P is less common but is predicted to have a strong impact on SF3B1 structure due to its location in an α-helix (Cretu et al., 2016). Similar to *sftb-1*[*K718E*], the two additional missense mutations, *sftb-1*[*R643C*] and *sftb-1*[*Q552P*] (**Figures S3a and S3b**), did not cause any evident phenotypes in *C. elegans*.

We were interested in producing animals with defective *sftb-1* function resulting in a viable and traceable phenotype for investigating modifiers of altered SFTB-1 activity. Since the effects of MDS-related missense mutations were reported to be additive in the yeast *SF3B1* ortholog Hsh155 (Carrocci et al., 2017), we generated three strains with each of the possible pairs of mutations: *sftb-1*[*R643C, K718E*]*, sftb-1*[*Q552P, K718E*]*, and sftb-1*[*Q552P, R643C*]. None of the three double mutants showed an obvious phenotype. However, animals simultaneously carrying the three missense mutations *sftb-1*[*Q552P, R643C, K718E*] displayed several temperature-sensitive developmental defects including partial sterility, protruding vulva, and developmental delay (Figure 1f–1h **and S3c–e**). These results suggest that each of the three missense mutations provokes a functional impact that becomes unmasked only when the three mutations are combined.

### Distinct *sftb-1* alleles produce alterations in alternative splicing and gene expression

The most commonly described AS defect caused by cancer-associated *SF3B1* mutations is the incorrect recognition of 3’ splice sites, producing aberrant transcripts (Darman et al., 2015; DeBoever et al., 2015; Pellagatti et al., 2018; Shiozawa et al., 2018; Wang et al., 2016) (Figure 2a). We investigated in *C. elegans* the transcriptional consequences of the K718E mutation, an equivalent of the most frequent *SF3B1* mutation K700E, by RNA-seq of two different strains with two independent K718E alleles (*cer3* and *cer7*) and their WT siblings at the L4 stage. Then, we used rMATS for robust detection of differential AS events between mutant and WT worms (Shen et al., 2014). Using both splice junction and exon body reads, our analysis detected 21985 AS events, of which 134 were significantly altered (FDR < 0.05) (**Table S1**). A more restricted cut-off (inclusion level difference > 0.1), as used in a previous rMATS analysis of *SF3B1*-mutated samples (Pellagatti et al., 2018), reduced the list of significant AS events to 78, with skipped exon (SE) being the most frequent event (Figures 2b **and** 2c).

**Figure 2.**
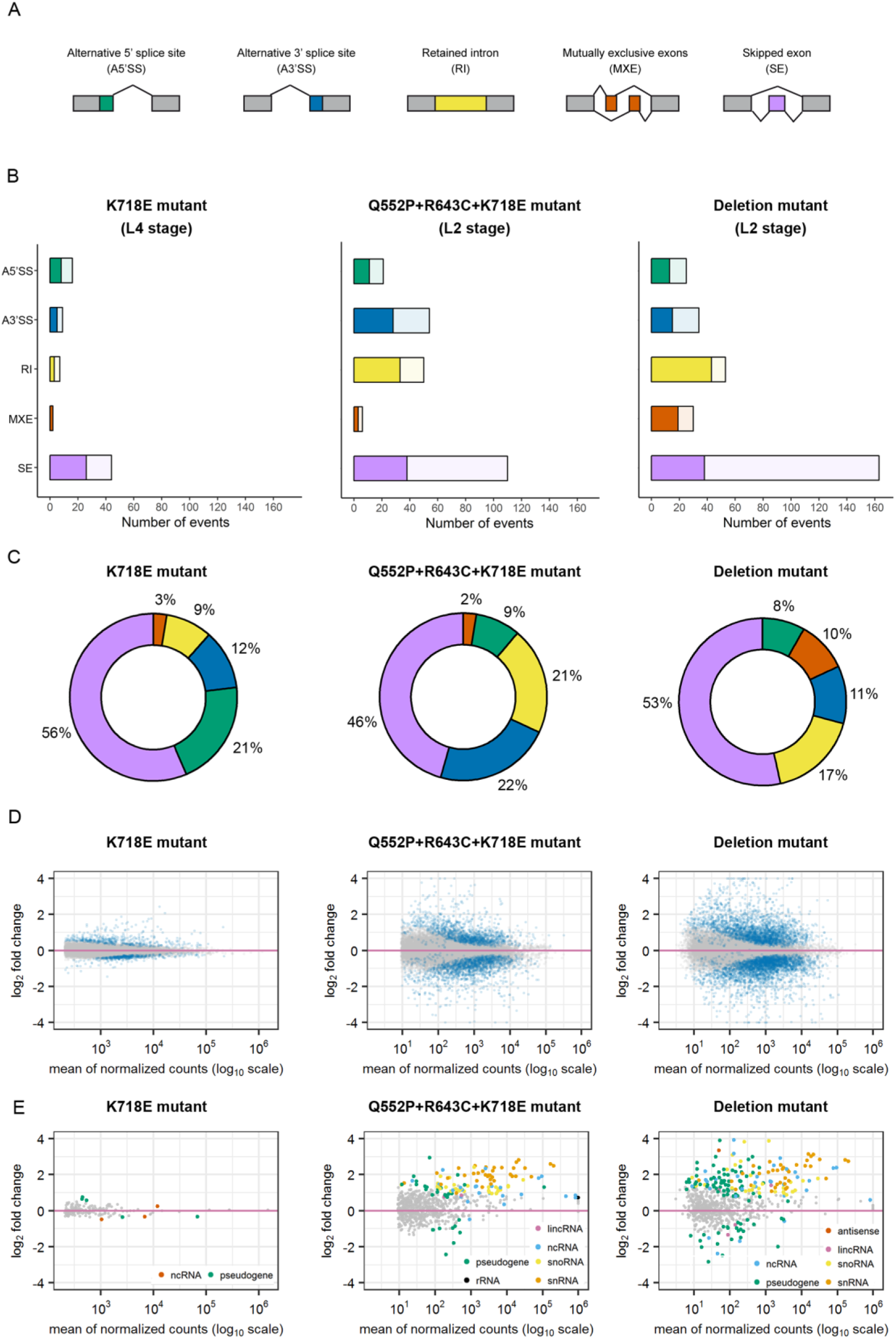
Different *sftb-1* mutations induce alternative splicing and gene expression alterations. (**A**) Types of AS events. Gray boxes represent constitutive exons, while colored boxes represent alternatively spliced exons. (**B**) Total number of significant (FDR < 0.05) AS alterations detected by rMATS with an inclusion level difference > 0.1. Event types are color coded as in panel A. Light colors represent AS events that were more significantly included in WT samples, while dark colors symbolize splicing alterations associated to mutant samples. (**C**) Doughnut charts showing the distribution of significant events represented in panel B by event type. In the three mutants, skipped exon (SE) was the most predominant alteration. (**D**) MA-plots displaying changes in protein-coding transcript expression levels (corrected log_2_fold change) over the mean of normalized counts, analyzed by DESeq2. Significant transcripts (padj < 0.05) are colored in blue. (**E**) MA-plots of expression changes in non-coding transcripts, represented as in panel D. Transcripts that are significantly deregulated are colored by transcript type. See also **Tables S1 and S2**.

Then, we investigated whether the two *sftb-1* alleles associated with a visible phenotype (the triple mutation allele *cer39*, and the deletion allele *cer6)* primarily induced cryptic 3’ splice site selection. It is worth noting that, in this experiment, larvae were harvested and processed for RNA isolation at the L2 stage to avoid possible indirect effects due to developmental defects observed after this stage. Both alleles were associated with a high number of deregulated AS events as determined by rMATS (Figure 2b), and the predominant alteration was also exon skipping (Figure 2c). We then explored if there were differentially expressed protein-coding (Figure 2d) and non-coding (Figure 2e) transcripts in the three RNA-seq datasets **(Table S2)**. As expected, the highest number of significantly altered transcripts was observed in the *cer6* allele.

In summary, we identified gene expression and alternative splicing alterations that are more frequent as *sftb-1* function becomes progressively compromised, with SE being the predominant AS defect.

### A subset of U2 snRNP and U2-associated components synthetically interacts with *sftb-1* mutants

In the search for selective vulnerabilities in *sftb-1* defective backgrounds, we hypothesized that *sftb-1* mutant animals would be more sensitive to knockdown of other splicing factors than their WT counterparts. Based on a previous *C. elegans* RNA interference (RNAi) collection of splicing factors (Kerins et al., 2010), we compiled and assayed a total of 104 RNAi clones targeting proteins that act in different steps of the splicing reaction. This led us to identify 27 splicing factors whose knockdown phenotype was enhanced in at least one of the three *sftb-1* single mutants, suggesting possible synthetic interactions (**Table S3**).

Among the 27 candidates, we selected 3 U2 snRNP components, including *sftb-1*, and 2 U2-associated proteins for further validation (Figure 3A). The identification of *sftb-1* as a candidate certified our approach since it demonstrated that any RNAi affecting *sftb-1*-related functions would have a stronger effect on *sftb-1* mutants. Since a partial loss-of-function mutant for the candidate *teg-4* was available, we proceeded to validate the interaction between *sftb-1* mutations and *teg-4*, the *C. elegans* ortholog of *SF3B3*, which physically interacts with SF3B1 HEAT repeats (Cretu et al., 2016). We found that the K718E mutation exacerbated the phenotype of *teg-4(oz210)* at 20°C (Mantina et al., 2009), whereas the Q552P mutation did so to a lesser extent (Figure 3B).

**Figure 3.**
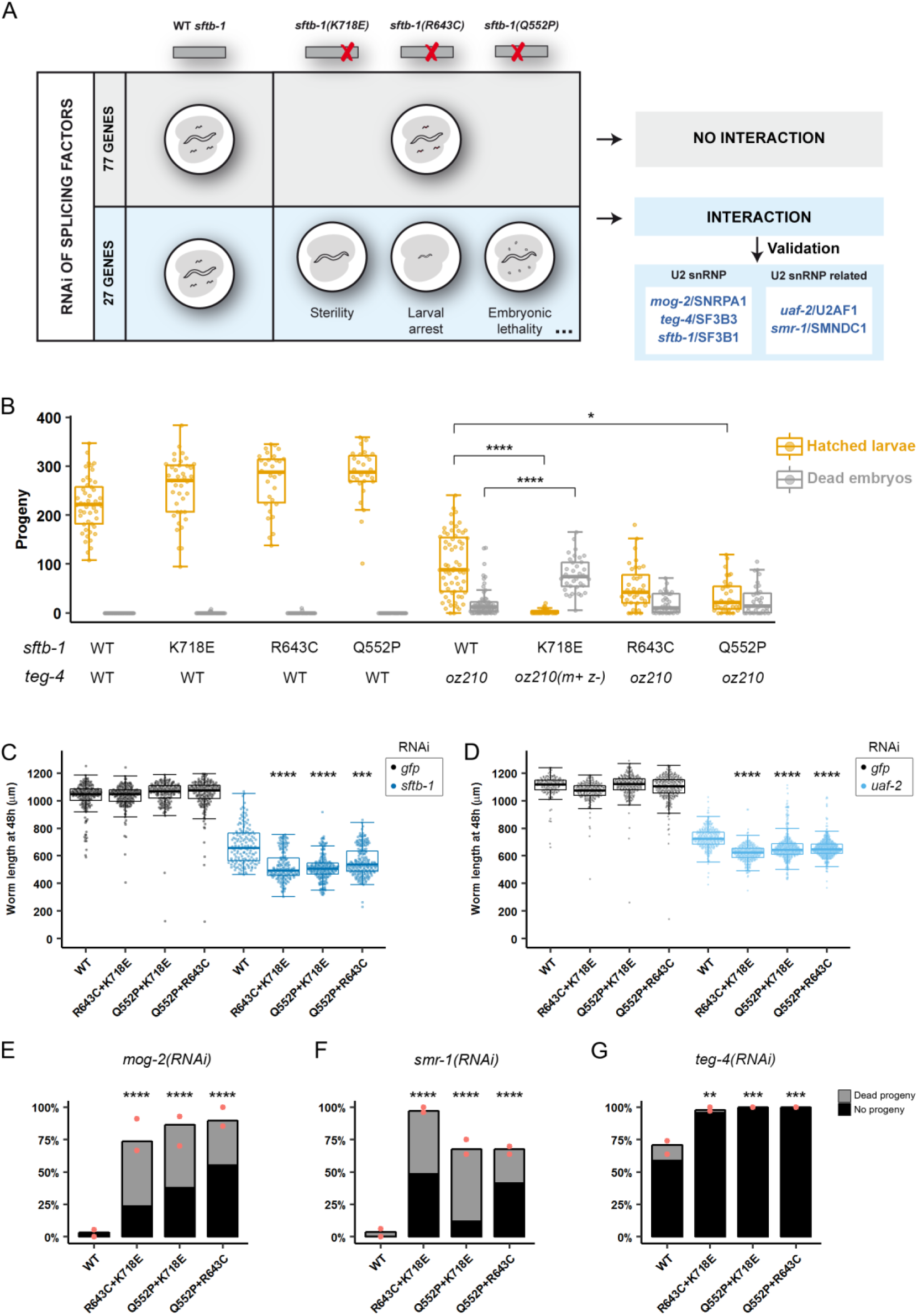
An RNAi screen targeting splicing genes leads to the validation of synthetic interactions between *sftb-1* mutants and five U2 snRNP and U2-associated components. (**A**) Schematics of the RNAi screen. In brief, WT or mutant worms bearing any of the three *sftb-1* single mutations were fed RNAi clones against a total of 104 splicing factors. 77 of these clones had the same effect in all genetic backgrounds. In contrast, we identified 27 clones that caused a stronger phenotype in at least one of the three single mutant strains compared to WT, which was indicative of a synthetic interaction. Listed are the 3 U2 snRNP and 2 U2 snRNP-related candidates from the screen that were selected for further validation. (**B**) *sftb-1[K718E]* and *sftb-1[Q552P]* synthetically interact with *teg-4(oz210)*. The number of hatched larvae (orange) and dead embryos (gray) laid by worms of the indicated genotype at 20°C is represented (n≥28; N=3). *m+ z-*, homozygous mutant progeny of heterozygous mothers. No significant differences were observed in the number of hatched larvae laid by *sftb-1[K718E], sftb-1[R643C]*, and *sftb-1[Q552P]* compared to WT. (**C**) Worm length after 48h of RNAi treatment against *gfp* (black) or *sftb-1* (blue) at 25°C (n≥150; N=2). (**D**) Worm length after 48h of RNAi treatment against *gfp* (black) or *uaf-2* (blue) at 25°C (n≥150; N=2). In both (c) and (d), worm length was not significantly different between *sftb-1* mutants and WT upon *gfp(RNAi)*. (**E-G**) Mean percentage of sterile worms observed upon RNAi of *mog-2* (E; n≥30), *smr-1* (F; n≥27) and *teg-4* (G; n≥34) at 25°C (N=2). Red dots indicate percent sterility observed in each replicate. Gray bars, P_0_-treated worms giving rise to <5 F_1_ larvae and some dead embryos (‘dead progeny’ category); black bars, P_0_-treated worms that laid neither larvae nor dead embryos (‘no progeny’ category). In (B), (C), and (D), data are shown as Tukey-style boxplots and overlaid dots representing measures in individual animals. Statistics: (B), (C), and (D), Kruskal-Wallis test with Dunn’s multiple comparisons test. Data were compared to *teg-4(oz210)* (B), or WT animals fed the same RNAi (C) and (D). (E), (F), and (G), Fisher’s exact test with Bonferroni correction for multiple comparisons. * p < 0.05, ** p < 0.01, *** p < 0.001, **** p< 0.0001. See also **Table S3**.

Then, we validated the rest of the candidates using RNAi. As we had observed that *sftb-1* mutations had an additive effect (Figure 1f–1h), we reasoned that *sftb-1* double mutants might be more sensitive to splicing perturbations than single mutants. Consistently, *sftb-1(RNAi)* led to an additive loss of function in double mutants, which arrested at earlier stages compared to WT worms (Figure 3c). A similar result was obtained with knockdown of *uaf-2*/*U2AF1* (Figure 3d), another hit from the RNAi screen.

Among the remaining U2 and U2-associated candidates, we selected *mog-2*/*SNRPA1* and *smr-1*/*SMNDC1* for further validation in the double-mutation backgrounds. Strikingly, whereas RNAi of *mog-2* resulted in very low penetrant sterility in WT worms, the incidence of the phenotype incremented to over 70% in the three *sftb-1* double-mutation strains (Figure 3e). A similar synthetic phenotype was observed upon knockdown of *smr-1* (Figure 3f). Finally, we further validated the synthetic interaction between *teg-4* and *sftb-1,* as *sftb-1* double mutants were more sensitive to *teg-4(RNAi)* (Figure 3g).

Together, our results suggest that targeting other splicing factors such as *SF3B3* to compromise their function can be an opportunity to selectively impair cancer cells harboring *SF3B1* mutations.

### The *C. elegans* mutations equivalent to human *U2AF1*^S34P^ and *SRSF2*^P95H^ genetically interact with *sftb-1*[*Q552P, R643C, K718E*] mutants

Along with *SF3B1* mutations, recurrent missense mutations in U2 small nuclear RNA auxiliary factor 1 (*U2AF1*) and serine/arginine-rich splicing factor 2 (*SRSF2*) were discovered in hematological malignancies (Yoshida et al., 2011), and later reported to be present in some solid tumors (Imielinski et al., 2012; Seiler et al., 2018b). Somatic hotspot mutations in these three splicing factors are only found in heterozygosis and are mutually exclusive, suggesting redundant functional impact or limited tolerance to splicing perturbations.

Missense mutations in *U2AF1* are commonly found at position S34, which corresponds to a residue located within the two zinc finger domains through which U2AF1 is thought to interact with the pre-mRNA (Ilagan et al., 2015). Similarly, *SRSF2* is typically mutated at position P95, mainly bearing missense mutations but also in-frame insertions or deletions (Yoshida et al., 2011).

Based on a recently reported synthetic interaction between *Srsf2*^P95H/+^ and *Sf3b1*^K700E/+^ in mice (Lee et al., 2018), and on the fact that *sftb-1* mutants were more sensitive to inactivation of some splicing factors (Figure 3), we wondered if the combination of *sftb-1*[*K718E*] with mutations equivalent to *U2AF1*^S34F^ or *SRSF2*^P95H^ would be tolerated in *C. elegans*.

Thus, we edited the worm orthologs of these genes to generate the *uaf-2*[*S42F*] and *rsp-4*[*P100H*] mutant strains. Neither of these two mutations caused any obvious phenotypes. We did not detect any interaction when combining each of these mutants with *sftb-1*[*K718E*] or the three *sftb-1* double-mutation strains. In contrast, *rsp-4*[*P100H*] and *uaf-2*[*S42F*] significantly increased the low percentage of sterile worms observed in the *sftb-1*[*Q552P, R643C, K718E*] background at 20°C (Figure 4a). The most dramatic effect was observed by the addition of the *uaf-2*[*S42F*] mutation to the *sftb-1* triple-mutation strain. Moreover, this combination of *uaf-2* and *sftb-1* mutations induced developmental delay at 20°C (Figure 4b). These results show for the first time in an animal model, a synthetic interaction between *sftb-1* and *uaf-2* missense mutations.

**Figure 4.**
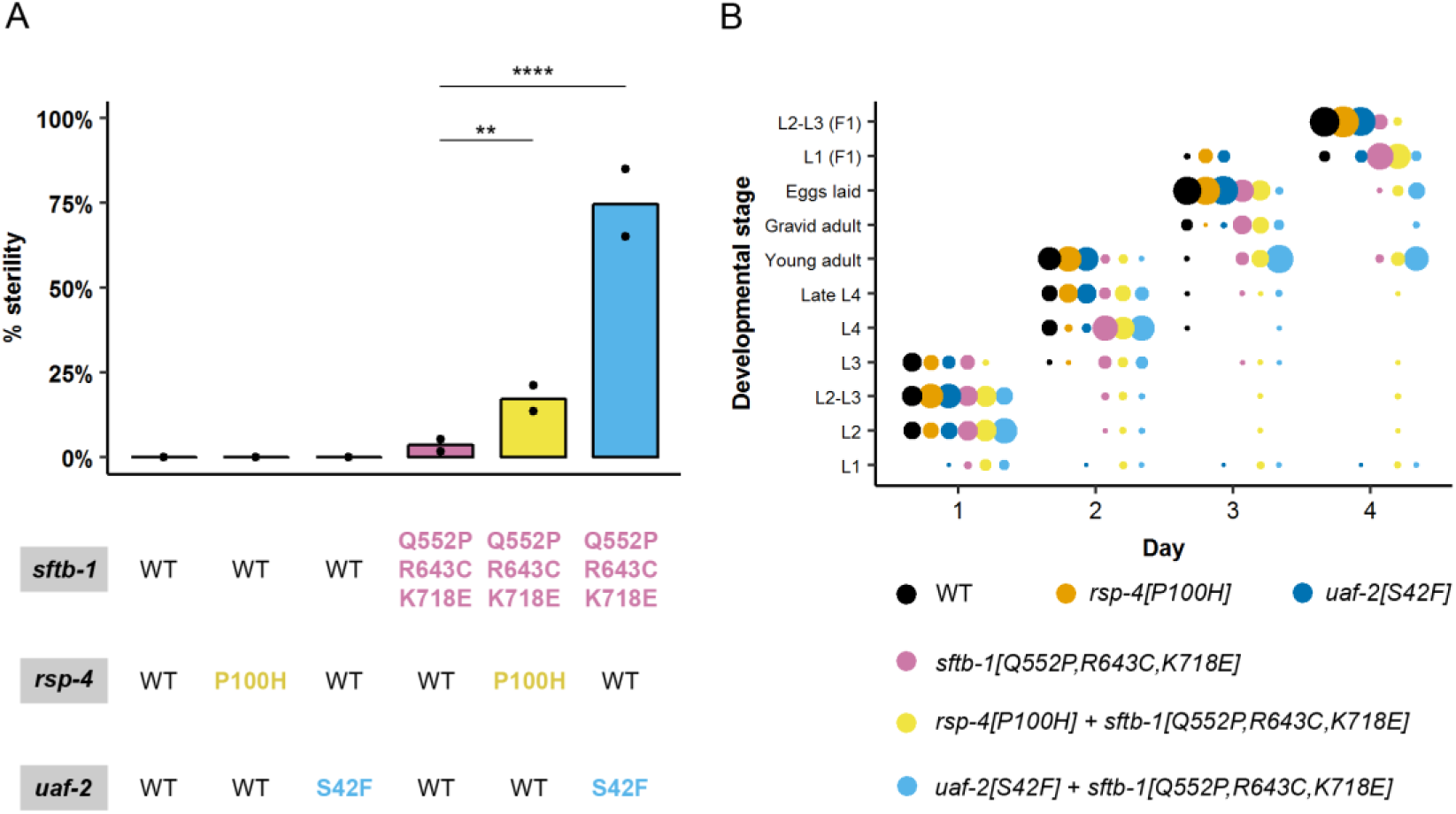
Genetic interaction between *rsp-4*/*SRSF2*, *uaf-2*/*U2AF1*, and *sftb-1* mutants. (**A**) Mean percentage of sterile worms of the indicated genotype at 20°C (N=2). Black dots denote observed values in each replicate. (**B**) *rsp-4[P100H]* and *uaf-2[S42F]* mutations gradually exacerbate the mild developmental delay displayed by the *sftb-1[Q552P, R643C, K718E]* mutant at 20°C. Dot sizes symbolize the percentage of worms at each developmental stage on any given day. In both panels, n≥110, N=2. Statistics: (A) Fisher’s exact test with Bonferroni correction for multiple comparisons. Lines indicate compared groups. ** p < 0.01, **** p < 0.0001.

### A humanized HEAT repeat 15 in SFTB-1 sensitizes *C. elegans* to pladienolide B

The SF3B1 C-terminal region contains 20 structural motifs known as HEAT repeats (HR), each one formed by two alpha helices joined by a short loop. These motifs display some structural plasticity and are involved in interactions with the pre-mRNA and other proteins (Cretu et al., 2016). Specific residues within SF3B1 HR15–17 configure the PB binding site (Cretu et al., 2018), and some amino acids in this region are not conserved in *C. elegans* (Figure 5A). In order to test if these sequence differences would affect the worms’ sensitivity to splicing inhibitors, we exposed WT animals and *sftb-1* cancer-related mutants to PB and to the synthetic molecule sudemycin D6 (Fan et al., 2011), which also targets SF3B1. Both drugs had no visible effects when delivered in liquid medium or via microinjection (**Table S4**). To reproduce the PB binding site in *sftb-1,* we used CRISPR to introduce four missense mutations (*cer144 allele*) that mimic the human HR15 (Figure 5B). The resulting strain was apparently wild type but sensitive to pladienolide B and insensitive to sudemycin D6. *sftb-1(cer144)* mutant larvae exposed to 10, 50, and 100 µM PB displayed a dose-dependent growth defect (Figure 5C).

**Figure 5.**
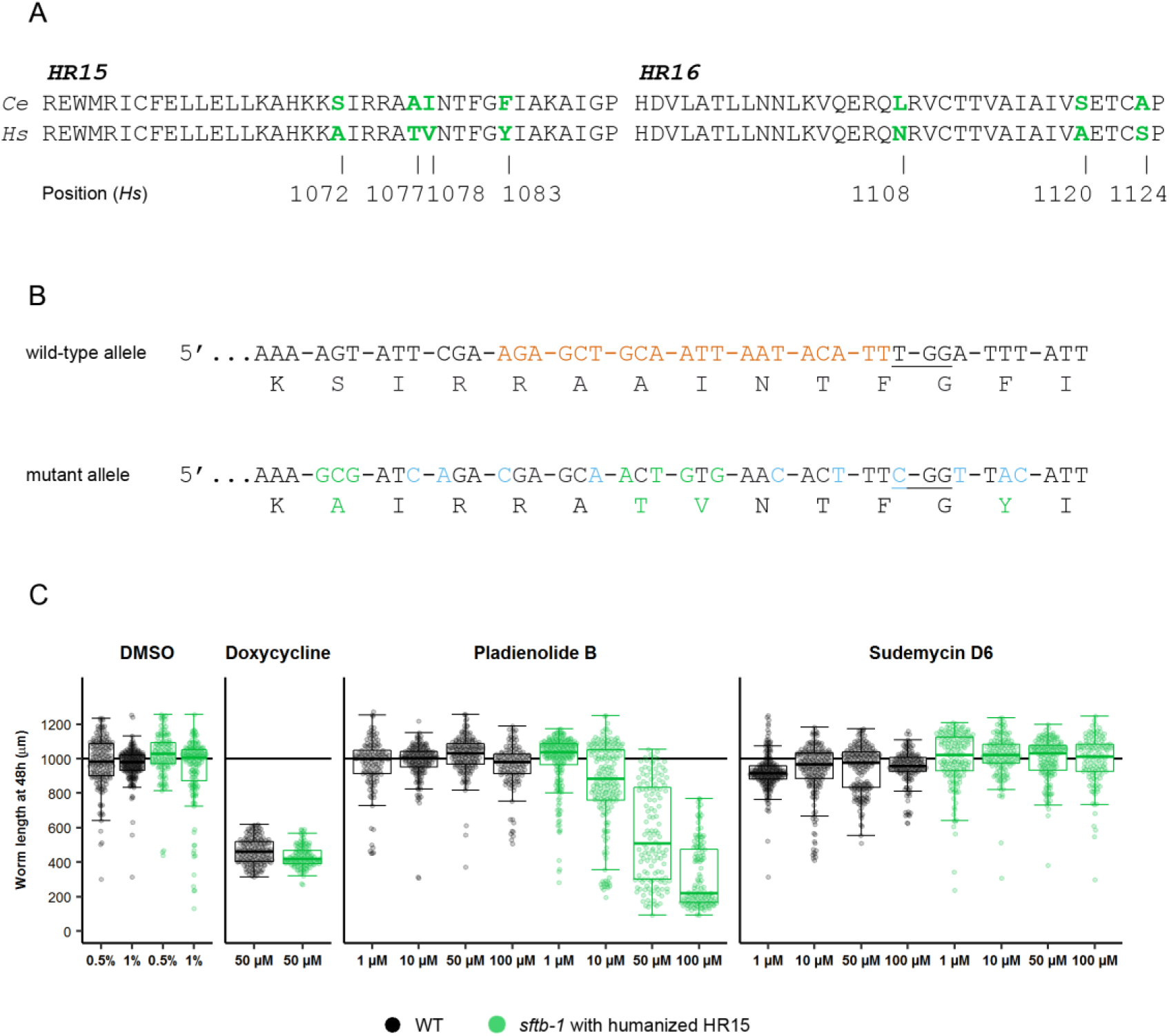
Humanization of the SFTB-1 HEAT repeat 15 confers sensitivity to Pladienolide B. (**A**) Protein sequence alignment of residues in HEAT repeats (HR) 15 and 16. Amino acids that differ in worm SFTB-1 (Ce) and human SF3B1 (Hs) are colored in green. Positions of these residues in the human sequence are indicated. (**B**) CRISPR/Cas9 design to “humanize” the SFTB-1 HR15. In the wild-type allele, the crRNA sequence used is indicated by orange nucleotides, and the protospacer-adjacent motif (PAM) is underlined. In the mutant (“humanized”) allele, synonymous mutations introduced to prevent partial recombination events and to improve primer specificity for mutant allele detection by PCR are indicated in blue. Missense mutations and the corresponding amino acids are colored in green. (**C**) Worm length after 48h of drug treatment at 25°C (n≥120; N=3). Doxycycline treatment induced larval arrest in both strains and was used as a positive control. Data are shown as Tukey-style boxplots and overlaid dots representing measures in individual animals. The horizontal line at 1000 µm indicates the expected body length of an adult worm. Body length of mutant worms treated with 10 µM and 50 µM PB differed statistically from that of mutant worms treated with 0.5% DMSO (p <0.0001). The same applied when comparing mutant worms treated with 100 µM PB and 1% DMSO. Significance was determined with the Kruskal-Wallis test with Dunn’s multiple comparisons test.

## DISCUSSION

*SF3B1* cancer-related mutations have been intensely studied in the past few years. They are particularly recurrent in MDS or UM, which commonly lack effective therapeutic approaches. In this study, we establish the first multicellular model for *SF3B1* mutations expressed non-conditionally from the endogenous locus. The rapid *C. elegans* life cycle (three days at 25°C) and the around 300 self-progeny obtained from a single hermaphrodite, among other features, make this model organism ideal for large-scale genetic and drug screens. The task of modeling human pathological mutations has been facilitated by the advent of the CRISPR/Cas9 genome editing technique, which has been efficiently implemented in this nematode.

As expected for a core splicing factor, and as previously reported in zebrafish (An and Henion, 2012) and mice (Isono et al., 2005), the *C. elegans SF3B1* ortholog is essential for development. Cancer cells bear *SF3B1* hotspot mutations in heterozygosis and depend on *SF3B1* WT function for viability. However, homozygous *sftb-1* missense mutations are viable in *C. elegans,* although they generate defects that are unmasked when additional alterations in the spliceosome are introduced. Such a feature is advantageous for investigating vulnerabilities of *SF3B1*-mutated tumors, which are not only conferred by missense mutations, but also by the loss of one *SF3B1* WT copy (Alsafadi et al., 2016; Paolella et al., 2017).

Differently from MDS with *SF3B1* mutations, where aberrant transcripts have been directly related to phenotypic traits (Dolatshad et al., 2015; Pellagatti et al., 2018), transcripts affected by *sftb-1*[*K718E*] do not produce any obvious phenotypes. Still, *Sf3b1*^K700E/+^ mice models only partially recapitulate the hematopoietic phenotype characteristic of MDS (Mupo et al., 2017; Obeng et al., 2016) and CLL (Yin et al., 2019). In these models, the splicing alterations caused by *Sf3b1*^K700E/+^ were similar to that reported in human tumors, but most of the affected transcripts were different, probably due to low conservation in intronic sequences.

RNA-seq of *sftb-1*[*K718E*] animals did not identify the aberrant use of alternative 3’ splice sites (A3’SS) as the predominant splicing defect caused by this mutation, as has been observed in human cells (Alsafadi et al., 2016; Darman et al., 2015; DeBoever et al., 2015; Pellagatti et al., 2018; Shiozawa et al., 2018). Indeed, in three distinct *sftb-1* mutants, we found a mild effect on AS, with SE being the most abundant splicing defect. Such disparity in the consequences of the mutation K700E/K718E between humans and nematodes may be explained by different factors. First, diverse computational pipelines have been used to assess differences in alternative splicing between WT and *SF3B1*-mutated samples, resulting in distinct conclusions. For instance, *Liberante et al.* reanalyzed previously published data and found that *SF3B1* mutation was mostly associated with differential skipped exon events, rather than A3’SS selection (Liberante et al., 2019). Second, intronic elements directing splicing have evolved differently in *C. elegans* compared to other organisms. *C. elegans* introns are typically shorter, do not have a consensus branch site (BS) sequence, and the vertebrate polypyrimidine (Py) tract is replaced by the consensus UUUUCAG/R sequence (Morton and Blumenthal, 2011). Third, only ∼35% of genes are alternatively spliced in *C. elegans* (Tourasse et al., 2017) compared to 95% in humans (Pan et al., 2008). Fourth, cell type-specific alternative isoforms generated by *sftb-1* mutations could have been underrepresented in our analyses as we sequenced the transcriptome of whole animals, and A3’SS selection can be regulated in a tissue-specific manner in *C. elegans* (Ragle et al., 2015). And fifth, the nonsense-mediated decay (NMD) system could efficiently eliminate transcripts with A3’SS resulting from *sftb-1* mutations, as occurs in human cells (Darman et al., 2015). However, we did not detect predominance of downregulation among the differentially expressed transcripts (Figures 2D **and** 2E).

Beyond its action on A3’SS usage, *SF3B1* missense mutations could have an impact on the structure, the interaction with other proteins in the spliceosome, or the robustness of the protein. In fact, the PB derivative H3B-8800, which induces lethality in spliceosome-mutant cancers by modulating the SF3b complex, affects AS activity similarly in WT and in *SF3B1*^K700E^ human cells (Seiler et al., 2018a). Therefore, *SF3B1*-mutated cells present vulnerabilities that can be exploited in synthetic lethal screens. We found that *sftb-1* mutants are sensitive to RNAi of different splicing factors, including itself, and we further validated the synthetic interaction with the inactivation of other components of the U2 snRNP. These results point to other components of the spliceosome as targets to preferentially kill cells with deficiencies in SF3B1 activity. Knowing the suitability of *sftb-1* mutants for RNAi screens, the collection of RNAi clones to be tested should be expanded, with particular interest placed on deubiquitinases and chromatin factors that have been related with SF3B1 functions (Kfir et al., 2015; Paolella et al., 2017).

Pladienolide and derivatives have been proven to be effective in selectively killing cells with *SF3B1* mutations (Obeng et al., 2016; Seiler et al., 2018a; Wu et al., 2018). We tested pladienolide B and sudemycin D6 in WT animals and *sftb-1* cancer-related mutants but did not observe any effects (**Table S4**). The inhibitory activity of these drugs is highly dependent on protein structure. By modifying four amino acids, we reproduced the human SF3B1 HR15 in *C. elegans* SFTB-1. As a result, worms became sensitive to PB but maintained resistance to sudemycin D6. Further humanization of SFTB-1 would expand the number of splicing inhibitors effective in *C. elegans*. It is worth mentioning that this is the first report on an invertebrate animal model sensitive to PB. Previously, splicing inhibition by PB *in vivo* was reported in yeast strains that were edited to humanize some Hsh155 HEAT repeats and mutated to block PB efflux (Hansen et al., 2019).

As an added value to the *C. elegans* toolkit, we have generated a strain that presents wild-type characteristics but sensitive to a splicing inhibitor. To date, chemical inhibition of splicing has not been possible in *C. elegans,* and therefore, this new PB-sensitive strain would facilitate splicing studies.

*SF3B1* mutations are mutually exclusive with other splicing factor mutations present in tumors. *U2AF1* and *SRSF2* missense mutations are reported in 11% and 12-15% of MDS, respectively (Xu et al., 2019), and very rarely overlap with *SF3B1* mutations, which are present in 17-28% of MDS patients (Lee et al., 2018; Seiler et al., 2018b). Hence cells may not handle all the AS defects produced by the combination of mutations in any of these three proteins. To further validate our model as a tool for identifying vulnerabilities in cells with compromised SF3B1 function, we mimicked *U2AF1*^S34F^ and *SRSF2*^P95H^ mutations in *C. elegans* and detected a functional interaction between both mutants and our *sftb-1* mutant carrying the three missense mutations. Additionally, *C. elegans rsp-4* and *uaf-2* missense mutations could be used for RNA-seq analyses and RNAi screens to study the consequences and vulnerabilities of these mutations, respectively.

We have demonstrated that *C. elegans* can be a valuable model to explore vulnerabilities of cells with mutations in *SF3B1*, although the molecular consequences of *sftb-1* mutations were different from the ones observed in other organisms. Replacement of several HEAT repeats in yeast Hsh155 with their human counterparts is functional and supports splicing (Carrocci et al., 2018). Similarly, partial or total replacement of *sftb-1* for its human ortholog could result in a more accurate model. To date, only one human gene has been functionally transplanted to *C. elegans* (McDiarmid et al., 2018), in part due to low efficiencies of inserting long DNA fragments by CRISPR. Such technical difficulties can now be bypassed by the use of Nested CRISPR (Vicencio et al., 2019). Thus, humanized *sftb-1* nematodes obtained by this method would be very efficient as *in vivo* models to further understand the impact of *SF3B1* mutations in cancer and as a platform for large-scale drug screens.

## Author Contributions

Conceptualization, X.S. and J.C.; Methodology, X.S., G.C., and J.C.; Validation, X.S.; Formal analysis, X.S. and A.E.C.; Investigation, X.S., D.K., H.B., and E.C.; Writing – original draft, X.S. and J.C., with help from the other authors; Writing – Review & Editing, X.S. and J.C.; Supervision, J.C.; Funding acquisition J.C.

## Acknowledgments

We acknowledge the members of the Cerón Laboratory for helpful discussions and comments on the manuscript. We also thank Alan Zahler and Sol Katzman for their advice in transcriptomic analyses. The *teg-4(oz210)* strain was kindly provided by Dave Hansen. Pladienolide B and Sudemycin D6 were gifts from Juan Valcárcel. This work has been supported by a grant from the Instituto de Salud Carlos III (ISCIII) to JC (PI15-00895), co-funded by FEDER funds/European Regional Development Fund (ERDF) — a way to build Europe. A.E.-C. is funded by ISCIII of the MINECO (reference PT17/0009/0019) and co-financed by FEDER. We thank CERCA Program / Generalitat de Catalunya for their institutional support. XS has an FPU PhD fellowship from MINECO and DK has an FI AGAUR fellowship from Generalitat de Catalunya.

## METHODS

### *Caenorhabditis elegans* strains

*C. elegans* strains were maintained using standard methods (Porta-de-la-Riva et al., 2012; Stiernagle, 2006) at the specified temperature. Before conducting the experiments, strains were grown for at least two generations at the experimental temperature. Unless otherwise specified, we used Bristol N2 as the WT strain. Mutant strains with single *sftb-1* missense mutations (CER218, CER220, CER236, and CER238) were outcrossed x2. WT siblings resulting from the second outcross were given specific strain names (CER217, CER221, CER237, and CER239, respectively) and kept for closer comparison. CER351, CER442 and CER450 strains were outcrossed x2. In order to maintain the *cer6* deletion, heterozygous mutants were crossed with the EG7893: *oxTi615 unc-119(ed3) III* strain so that a red fluorescent marker was inserted 0.15 cM away from the *sftb-1* WT locus.

### Protein alignments

*H. sapiens* SF3B1 (Uniprot: O75533) and *C. elegans* SFTB-1 (Uniprot: G5EEQ8) protein sequences were aligned using the T-Coffee algorithm (Notredame et al., 2000) with default settings. **Figure S1** was generated with Jalview (Waterhouse et al., 2009).

### CRISPR/Cas9 mutant and reporter strains

Specific guide RNAs were designed using both *Benchling* (www.benchling.com) and *CCTop* (Stemmer et al., 2015) online tools. All CRISPR/Cas9 mutant and reporter strains were obtained following a co-CRISPR strategy (Kim et al., 2014) using *dpy-10* as a marker to enrich for genome-editing events. In all cases, mixes were injected into gonads of young adult P_0_ hermaphrodites using the XenoWorks Microinjection System and following standard *C. elegans* microinjection techniques. F_1_ progeny was screened by PCR using specific primers and F_2_ homozygotes were confirmed by Sanger sequencing. As we optimized the protocol for CRISPR/Cas9 genome editing in our laboratory while preparing this manuscript, different reagents and conditions were used to produce the distinct mutants. The *sftb-1(cer6)* deletion mutant allele was obtained by nonspecific double-strand break repair on our first attempt to generate the K718E mutation. Only in this experiment, guide RNAs and Cas9 were expressed from plasmids that were purified with the NucleoBond® Xtra Midi Kit (Macherey-Nagel). In the remaining mixes, ribonucleoprotein complexes containing crRNA, tracrRNA and Cas9 were annealed at 37°C for 15 minutes prior to injection. All the reagents and injection mixes used in this study are listed in **Table S5**.

### *C. elegans* microscopy

Worms were mounted on 2% agar pads with 10 mM tetramisole hydrochloride and imaged using a ZEISS Axio Observer Z1 inverted fluorescence microscope. Images were processed using ZEISS ZEN 2012 (blue edition) software and Fiji.

### Gonad dissection and DAPI staining

Worms expressing *mCherry::SFTB-1* were anesthetized with 0.33 mM tetramisole hydrochloride and their heads cut using a pair of 25-gauge needles to extrude gonad arms. Released gonads were fixed in 4% paraformaldehyde in PBS for 20–30 minutes and washed with PBS + 0.1% Tween 20 for 10 minutes. After performing 3 washes, gonads were transferred to a microscope slide and mounted using 10 µl DAPI Fluoromount-G® (SouthernBiotech). Images were acquired the same day to avoid mCherry signal loss.

### *sftb-1* deletion mutant larval arrest

Synchronized L1-arrested F_2_ worms from heterozygous *sftb-1(cer6)/oxTi615* P_0_s were seeded onto NGM plates with OP50 and imaged after 48 hours of growth at 20°C using a ZEISS Axio Observer Z1 inverted fluorescence microscope at 10X magnification. Body length was measured manually by drawing the midline of the worm using the freehand line tool in Fiji.

### Phenotypic characterization

Similar experiments were conducted to characterize *sftb-1[Q552P, R643C, K718E], rsp-4[P100H]* and *uaf-2[S42F]* mutant phenotypes. Briefly, synchronized L1-arrested worms of the specified genotype were first seeded onto NGM plates with OP50 and, after a few hours to allow recovery from the L1 arrest, 60 animals per genotype were separated individually to 12-well NGM plates seeded with OP50. Plates were incubated at 20°C (Figures 4 **and S3c–E**) or 25°C (Figure 1f–h). The developmental stage of individual worms was assessed visually every 24 hours under a stereomicroscope based on animal size and germline morphology. Phenotypes were scored at day 4. In order to determine brood sizes of *sftb-1[Q552P, R643C, K718E]* and WT animals, a proportion of worms were transferred into new 35-mm NGM plates at day 4 and subsequently moved to fresh plates every 12–24 h until egg laying ceased. Progeny were counted 36–48 h after passages.

### RNA isolation

To study the transcriptional consequences of the K718E mutation, we used two homozygous mutant lines (CER218 and CER220) derived from two independent CRISPR/Cas9 events (*cer3* and *cer7* alleles, respectively), and the corresponding WT siblings (CER217 and CER221). Synchronized L4 animals were collected in M9 after growing for 41 hours at 20°C and total RNA was isolated using TRI Reagent (MRC, Inc.) and the PureLink® RNA Mini Kit (Ambion), following the manufacturer’s instructions. *sftb-1(cer6)* homozygous larvae were obtained by automatically sorting non-red animals from a synchronized CER191 population grown at 20°C for 23 hours, using the COPAS Biosorter system (Union Biometrica). We conducted two biological replicates with an average of 1.6% non-red recombinants and 5.25% red contaminants in the samples. The same size and fluorescence window was used to sort two N2 WT populations collected under the same experimental conditions. Two CER276 biological replicates were collected independently without sorting, after growing for 23 hours at 20°C. Total RNA from the six samples was extracted using TRI Reagent, followed by aqueous and organic phase separation using 1-Bromo-3-chloropropane (Sigma-Aldrich), and RNA was precipitated with 2-propanol. After treatment with TURBO DNase (Thermo Fisher Scientific), RNA was purified following a phenol/chloroform extraction protocol.

In all cases, total RNA was quantified by Qubit® RNA BR Assay kit (Thermo Fisher Scientific) and the integrity was checked by using the Agilent RNA 6000 Nano Kit (Agilent).

### RNA sequencing

Four RNA-seq libraries (from CER217, CER218, CER220, and CER221 strains) were prepared with the KAPA Stranded mRNA-Seq Illumina® Platforms Kit (Roche-Kapa Biosystems) following the manufacturer’s recommendations. Briefly, 500 ng of total RNA was used as the input material, the poly-A fraction was enriched with oligo-dT magnetic beads and the mRNA was fragmented. Strand specificity was achieved during the second strand synthesis performed in the presence of dUTP instead of dTTP. The blunt-ended double stranded cDNA was 3’adenylated and Illumina indexed adapters from TruSeq™ Stranded Total RNA Sample Preparation Kit with Ribo-Zero Gold (Illumina) were ligated.

The remaining six samples (two biological replicates from N2, CER276, and homozygous *sftb-1(cer6)* larvae) were prepared using the TruSeq™ Stranded Total RNA kit protocol (Illumina). Briefly, rRNA was depleted from 500 ng of total RNA using the Ribo-Zero Gold rRNA Removal Kit and fragmented by divalent cations, with a major peak at 160 nt. Following the fragmentation, first and second strand synthesis was performed, also ensuring the library directionality. The cDNA was adenylated and ligated to xGen® Dual Index UMI Adapters (IDT) for paired-end sequencing.

Both types of libraries ligation products were enriched with 15 PCR cycles and the final library was validated on an Agilent 2100 Bioanalyzer with the Agilent DNA 1000 Kit (Agilent).

The libraries were sequenced on HiSeq 2000 (Illumina) in paired-end mode with a read length of 2×76bp using the HiSeq SBS Kit V4 50 cycle kit in a fraction of a sequencing v4 flow cell lane (HiSeq PE Cluster Kit V4 - cBot). Image analysis, base calling, and quality scoring of the run were processed using the manufacturer’s software Real Time Analysis (RTA 1.18.66.3) and followed by generation of FASTQ sequence files.

### Bioinformatic analysis

RNA-seq reads were mapped against *C. elegans* reference assembly (WBcel235.87) with STAR (Dobin et al., 2013). Genes and transcripts were quantified with RSEM (Li and Dewey, 2011). Differential expression analysis was performed with DESeq2 R package (Love et al., 2014), and detection of differential alternative splicing events was done with rMATS (Shen et al., 2014).

### RNAi screen

Information about the selected splicing genes was collected from SpliceosomeDB (Cvitkovic and Jurica, 2013) or from the literature. RNAi clones were obtained from the ORFeome library (Rual et al., 2004) or the Ahringer library (Kamath et al., 2003) and clone insert size was validated by PCR. For screening, we used 24-well plates containing NGM supplemented with 50 µg/mL ampicillin, 12.5 µg/mL tetracycline and 3 mM IPTG. Each well was seeded with 80 µl of bacterial culture and dsRNA expression was induced overnight at room temperature. WT and mutant worms were tested in duplicates. Between 10 and 20 worms synchronized at the L1 stage were seeded in each well, and phenotypes were scored visually at the adult stage (72–96 h post-seeding at 25°C). Bacteria expressing dsRNA against *gfp* were used as negative control.

### Interaction between *teg-4* and *sftb-1* mutations

Worms of the indicated genotype were singled onto individual 35-mm NGM plates with OP50 at the L4 stage and transferred to new plates every 12 hours until all eggs were laid. Numbers of dead embryos and hatched larvae were scored 24 and 48 hours after passages, respectively. Since the double mutant *sftb-1(K718E); teg-4(oz210)* could not be maintained in homozygosis, F_1_ worms from *sftb-1(K718E); teg-4(oz210/+)* P_0_ worms, including animals homozygous and heterozygous for *teg-4(oz210)*, were picked at the L4 stage and their progeny scored. These F_1_s were genotyped after conducting the experiment to include only *sftb-1(K718E)* and *sftb-1(K718E); teg-4(oz210)* data.

### Validation of candidates from the RNAi screen

In all cases, we used 55-mm or 35-mm NGM plates supplemented with 50 µg/mL ampicillin, 12.5 µg/mL tetracycline and 3 mM IPTG, seeded with specific RNAi bacterial clones. RNAi targeting *gfp* was used as negative control.

In Figures 3c **and** 3d, synchronized L1-arrested worms of the indicated genotypes were seeded onto 55-mm plates, and imaged directly on the plates after 48 hours of growth at 25°C using a stereomicroscope. Body length was determined automatically with WormSizer (Moore et al., 2013).

In Figures 3e–g, synchronized L1-arrested animals were seeded onto 55-mm plates and separated into individual 35-mm plates at the L4 stage. Worms were transferred every 24 hours and the presence of dead embryos was determined 24 hours after passages.

### Number of germ cells assay

WT or *sftb-1(cer6)* homozygous worms were collected at the indicated time points, fixed using Carnoy’s solution (60% absolute ethanol, 30% chloroform, 10% acetic acid), and washed three times in PBS + 0.1% Tween 20. Whole worms were transferred to a microscope slide and stained with DAPI Fluoromount-G® in order to visualize and count germ cell nuclei.

### Spliceosome inhibitors experiments

Experiments in liquid media were conducted in 96-well plates. Experiments in Table S4 were conducted as follows: 25 µl of dead OP50 resuspended in a mix of freshly prepared S Medium (Stiernagle, 2006), 4 µg/ml cholesterol, 250 µg/ml streptomycin, and 62.5 µg/ml tetracycline were added to each well.

Subsequently, 25 µl 2.4X drug and 10 µl L1-arrested worms at a density of 1–2 worms/µl were added to each well. Each condition was tested in duplicate in the same 96-well plate. Plates were incubated in a humid chamber with gentle agitation at room temperature for the indicated time period. 25 µl water (experiments 1–3) or 0.5% DMSO (Sigma-Aldrich) (experiments 4 and 6) were used as negative controls. In experiment 5, 50 µM udemycin D6 or 0.5% DMSO were injected into the gonads or intestine of young adult hermaphrodites using standard microinjection techniques, and injected animals were recovered in individual 35-mm NGM plates with OP50 and incubated at 20°C.

The protocol for liquid experiments in Figure 5C was slightly modified. In brief, 64.5 μl S Medium supplemented with 5 μg/ml cholesterol, 50 μg/ml streptomycin, and 50 μg/ml ampicillin were added to each well. This supplemented S-Medium was used to resuspend a pellet of dead OP50 to an optical density of 1.1-1.2, and 25 μl of bacteria were dispensed to each well. Subsequently, 10 μl L1-arrested worms at a density of 4–6 worms/μl, and 0.5 μl 200X drug were added to each well. Each condition was tested in triplicate in the same 96-well plate. Plates were incubated in a humid chamber at 25°C.

### Statistical analyses

In all figure legends, ‘N’ denotes the number of independent replicate experiments performed, while ‘n’ indicates the total number of animals analyzed in each condition (different genotypes are separated by commas). Statistical analyses were performed in GraphPad Prism 6 and R. Statistical tests used are reported in the figure legends. Figures 1a, 1d, 1f–h, 2b–e, 3b–g, 4, 5c, **S2b**, and **S3c–e** were generated by using the ggplot2 R package.

### Data availability

The RNA sequencing data generated in this study have been deposited in the Gene Expression Omnibus repository and are available under the accession number GEO: GSE129642.

## Supplementary information

**Figure S1.** Most of the SF3B1 mutated residues are conserved in the *C. elegans* ortholog *sftb-1*.

**Figure S2.** *sftb-1(cer6)* deletion mutant.

**Figure S3.** *sftb-1* missense mutations design and *sftb-1[Q552P, R643C, K718E]* phenotypes at 20°C.

**Table S1.** Alternative splicing analysis of *sftb-1* mutants by rMATS.

**Table S2.** Differential expression analysis of *sftb-1* mutants by DeSeq2.

**Table S3.** Information and phenotypes of the RNAi splicing screen.

**Table S4.** First round of experiments with spliceosome inhibitors.

**Table S5.** CRISPR reagents and injection mixes.

