## Supplementary material for "CRISPR editing of *sftb-1*/SF3B1 in *C. elegans* allows the identification of synthetic interactions with cancer-related mutations and the chemical inhibition of splicing": Figures S1, S2, S3 (from another version of this preprint: version 3, 2019-05-15)

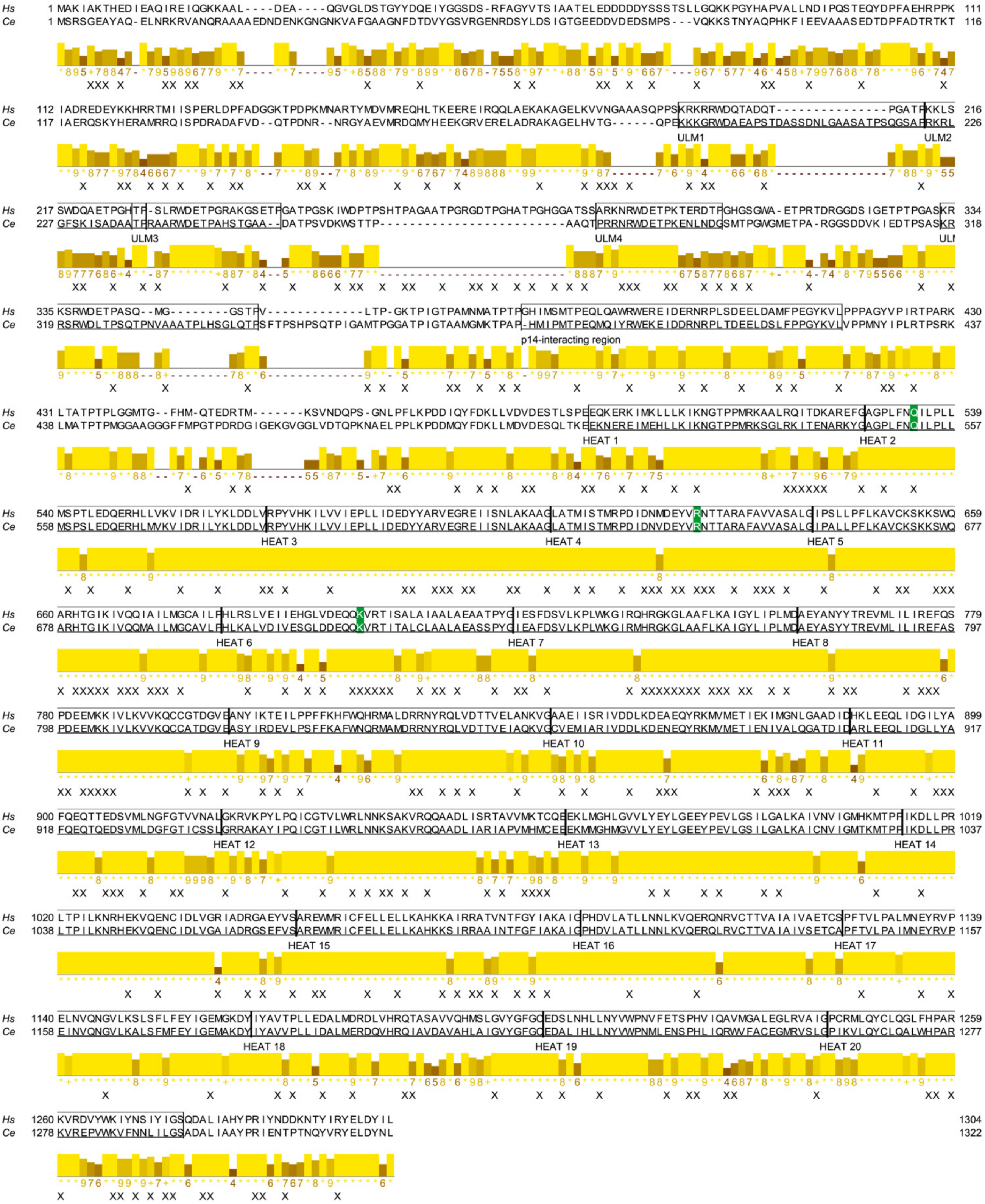

Hs
Ce
ULM1
ULM2
ULM3
ULM4
p14-interacting region
HEAT 1
HEAT 2
HEAT 3
HEAT 4
HEAT 5
HEAT 6
HEAT 7
HEAT 8
HEAT 9
HEAT 10
HEAT 11
HEAT 12
HEAT 13
HEAT 14
HEAT 15
HEAT 16
HEAT 17
HEAT 18
HEAT 19
HEAT 20

**Figure S1. Most of the *SF3B1* mutated residues are conserved in the *C. elegans* ortholog *sftb-1.* Related to Figure 1.**

Protein sequence alignment of human SF3B1 *(Hs)* and worm *sftb-1 (Ce).* The yellow histogram represents the degree of conservation in terms of amino acid physico-chemical properties. The numerical index below reflects conservation in a quantitative manner, where * corresponds to the maximum degree of conservation. U2AF binding motifs (ULMs), p14-interacting region and HEAT domains are indicated in boxes. A cross symbol (X) below the conservation score indicates positions where missense mutations have been reported in human cancers (COSMIC database) at different frequencies. Residues that are relevant for this study are highlighted in green.

A

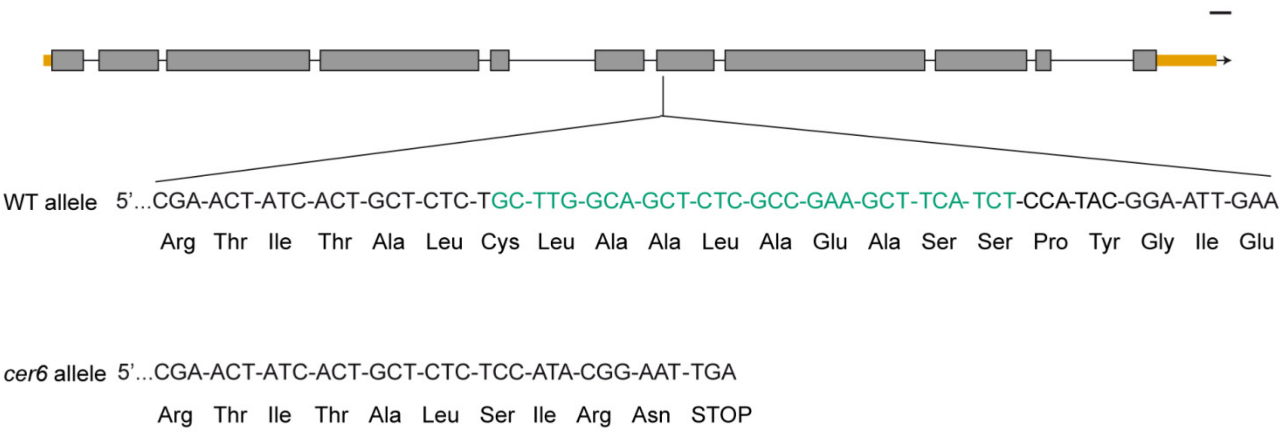


B

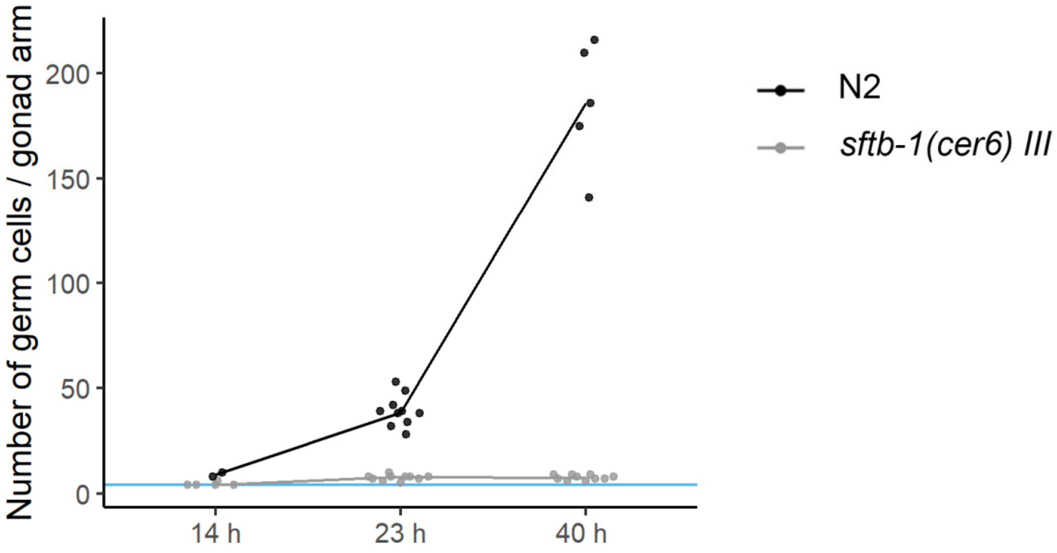


**Figure S2. *sftb-1(cer6)* deletion mutant. Related to Figure 1.**

(A) Schematic representation of the *sftb-1* locus. Gray boxes indicate exons, connecting lines indicate introns. Left and right orange boxes represent predicted 5' and 3' untranslated regions, respectively. In green, 29 nucleotides that are deleted in the *cer6* allele. Predicted amino acids resulting from translating the WT or *cer6* alleles are shown below the DNA sequence. Scale bar, 100 nucleotides.

(B) Germ cell number of WT or homozygous *sftb-1(cer6)* synchronized larvae at 14, 23, and 40 hours post-L1. In WT worms, differentiating spermatocytes were not included at the last time point, and the number of germ cells was counted in one gonad arm only. Black and gray lines connect the median at each time point in WT and *sftb-1(cer6)* worms, respectively. The blue line represents the four germ cells that are normally present in a recently hatched L1 larva, and that will resume proliferation during the L1 stage.

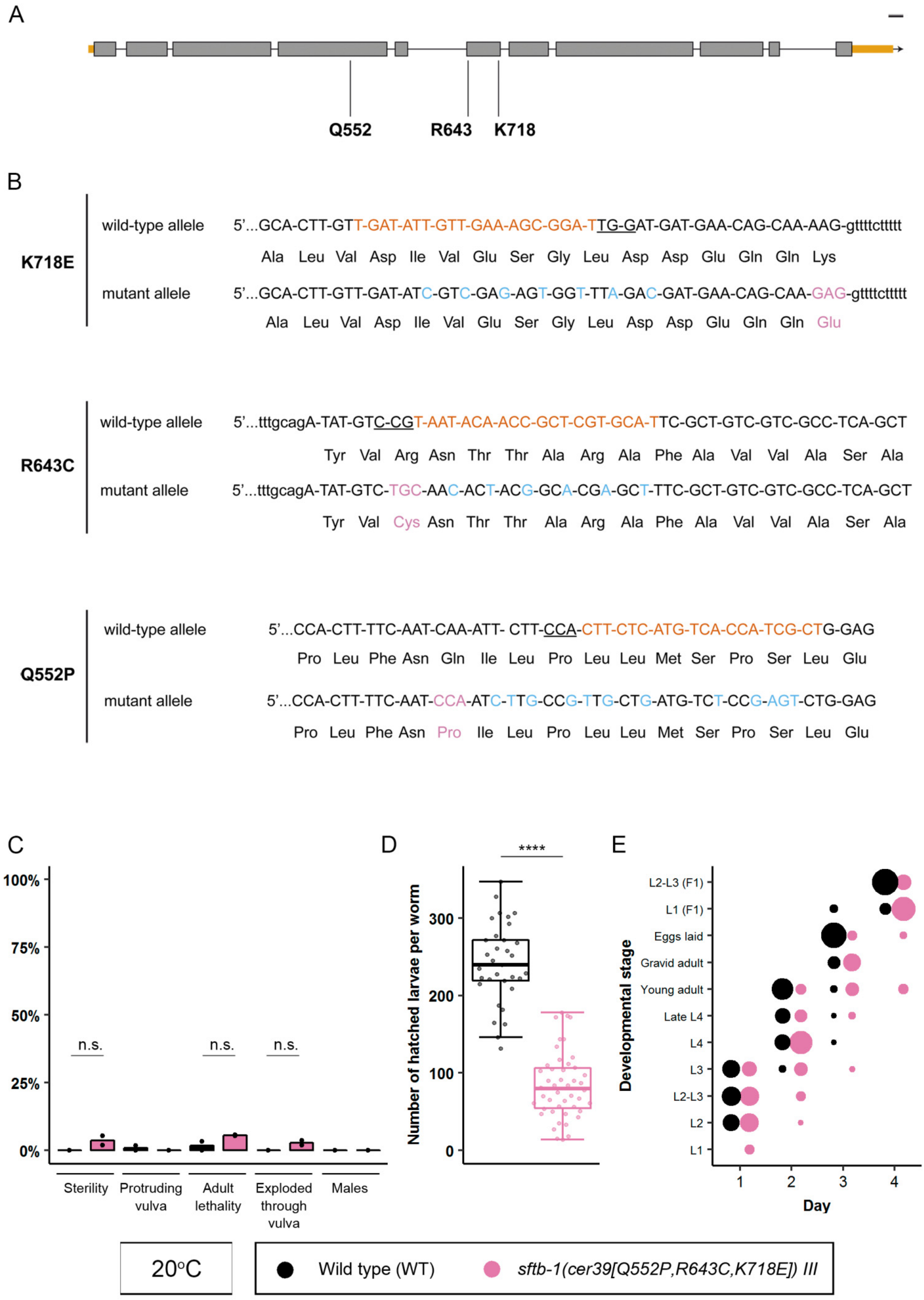
A
Q552
R643
K718
B
K718E
wild-type allele
5'...GCA-CTT-GTT-GAT-ATT-GTT-GAA-AGC-GGA-TTG-GAT-GAT-GAA-CAG-CAA-AAG-gttttcttttt
Ala Leu Val Asp Ile Val Glu Ser Gly Leu Asp Asp Glu Gln Gln Lys
mutant allele
5'...GCA-CTT-GTT-GAT-ATC-GTC-GAG-AGT-GGT-TTA-GAC-GAT-GAA-CAG-CAA-GAG-gttttcttttt
Ala Leu Val Asp Ile Val Glu Ser Gly Leu Asp Asp Glu Gln Gln Glu
R643C
wild-type allele
5'...tttgcagA-TAT-GTC-CGT-AAT-ACA-ACC-GCT-CGT-GCA-TTC-GCT-GTC-GTC-GCC-TCA-GCT
Tyr Val Arg Asn Thr Thr Ala Arg Ala Phe Ala Val Val Ala Ser Ala
mutant allele
5'...tttgcagA-TAT-GTC-TGC-AAC-ACT-ACG-GCA-CGA-GCT-TTC-GCT-GTC-GTC-GCC-TCA-GCT
Tyr Val Cys Asn Thr Thr Ala Arg Ala Phe Ala Val Val Ala Ser Ala
Q552P
wild-type allele
5'...CCA-CTT-TTC-AAT-CAA-ATT- CTT-CCA-CTT-CTC-ATG-TCA-CCA-TCG-CTG-GAG
Pro Leu Phe Asn Gln Ile Leu Pro Leu Leu Met Ser Pro Ser Leu Glu
mutant allele
5'...CCA-CTT-TTC-AAT-CCA-ATC-TTG-CCG-TTG-CTG-ATG-TCT-CCG-AGT-CTG-GAG
Pro Leu Phe Asn Pro Ile Leu Pro Leu Leu Met Ser Pro Ser Leu Glu
C
100%
75%
50%
25%
0%
n.s.
n.s.
n.s.
Sterility
Protruding vulva
Adult lethality
Exploded through vulva
Males
D
****
Number of hatched larvae per worm
300
200
100
0
E
L2-L3 (F1)
L1 (F1)
Eggs laid
Gravid adult
Young adult
Late L4
L4
L3
L2-L3
L2
L1
Developmental stage
1
2
3
4
Day
20°C
Wild type (WT)
sftb-1(cer39[Q552P,R643C,K718E]) III

**Figure S3. *sftb-1* missense mutations design and *sftb-1[Q552P, R643C, K718E]* phenotypes at 20°C. Related to Figure 1.**

(A) Schematic representation of the *sftb-1* locus, as in **Figure S2**. The position of the triplets encoding the three mutated residues in this study (Q552, R643 and K718) is indicated. Scale bar, 100 nucleotides.

(B) Molecular details of the three *sftb-1* missense mutations generated by CRISPR/Cas9. Lowercase letters indicate introns, whereas uppercase letters depict exonic sequences. In WT alleles, the crRNA sequence used is indicated by orange nucleotides, and the protospacer-adjacent motif (PAM) is underlined. In mutant alleles, synonymous mutations introduced to prevent partial recombination events and to improve primer specificity for mutant allele detection by PCR are indicated in blue. Missense mutations and the corresponding amino acids are colored in pink.

(C) Bars represent mean incidence of different phenotypes observed in *sftb-1[Q552P, R643C, K718E]* worms at 20°C, while dots represent percentages observed in each replicate (n=116, 109; N=2).

(D) Total number of progeny laid by WT or *sftb-1[Q552P, R643C, K718E]* worms at 20°C (n=33, 46; N=2). Dots represent values for each individual animal, overlaid to Tukey-style boxplots.

(E) *sftb-1[Q552P, R643C, K718E]* animals present a mild developmental delay at 20°C. Dot sizes represent the proportion of the population at each stage (n=116, 109; N=2)

Statistics: (C), Fisher's exact test (D), Student's *t*-test (unpaired, two-tailed). n.s., no significant difference, **** $p < 0.0001$
